# C99 selectively accumulates in vulnerable neurons in Alzheimer’s disease

**DOI:** 10.1101/527572

**Authors:** Maria V. Pulina, Maya Hopkins, Vahram Haroutunian, Paul Greengard, Victor Bustos

## Abstract

**Introduction:** The levels and distribution of amyloid deposits in the brain does not correlate well with Alzheimer’s disease (AD) progression. Therefore, it is likely that Amyloid-precursor-protein proteolytic fragments other than beta-amyloid contribute to the onset of AD.

**Methods:** We developed a sensitive assay adapted to the detection of C99, the direct precursor of beta-amyloid. Three postmortem groups were studied: control with normal and stable cognition; subjects with moderate AD, and individuals with severe AD. The amount of C99 and beta-amyloid was quantified and correlated with the severity of AD.

**Results:** C99 accumulates in vulnerable neurons, and its levels correlate with the degree of cognitive impairment in patients suffering from AD. In contrast, beta-amyloid levels are increased in both vulnerable and resistant brain areas.

**Discussion:** These results raise the possibility that C99, rather than beta-amyloid plaques, is responsible for the death of nerve cells in Alzheimer’s disease.

## 1. Introduction

Alzheimer’s disease (AD) is the most common type of dementia, characterized by accumulation of extracellular and intracellular beta-amyloid aggregates, intracellular paired helical fragments and extensive age-dependent neuronal loss in vulnerable areas [1-3]. Robust genetic and biochemical evidence points to alterations of Amyloid-precursor protein (APP) metabolism as causative in the development of Alzheimer’s disease. In the amyloidogenic pathway, APP is cleaved by beta-secretase (BACE) to generate sAPPb and C99. Gamma-secretase then cleaves C99 to generate beta-amyloid and AICD. Accumulation of beta-amyloid is a cardinal feature of Alzheimer’s disease, but its role in causing injury in AD remains unclear. Thus, the distribution of beta-amyloid deposits in AD brains has a poor correlation with dementia severity, loss of neural function and cognitive impairment [4-9]. Therefore, it is possible that APP metabolites other than beta-amyloid make a significant contribution to AD pathophysiology. In addressing this issue, a challenge has been the inability to determine the distribution and levels of specific APP metabolites in intact cells and tissues. We have approached this problem by adapting the proximity ligation assay (PLA) for the detection of C99. In this assay, two primary antibodies recognize the target epitopes on the protein of interest. If the epitopes are in close proximity, secondary antibodies covalently bound to complementary DNA strands participate in rolling circle DNA synthesis. The DNA synthesis reaction results in a one thousand-fold amplification of the DNA circle. Fluorescently-labeled complementary oligonucleotide probes added to the reaction bind to the amplified DNA. The resulting fluorescent signal can be detected by fluorescence microscopy. This technique offers great sensitivity, since powerful rolling circle amplification is used. It also offers great selectivity, since dual target recognition is used [10, 11]. Here, we used antibodies generated against both the N and C terminal domains of C99 in cultured cells and found that we can distinguish C99 from beta-amyloid, full-length APP, C83, and AICD. The adaptation of the proximity ligation assay to the detection of C99 provides a general framework for the detection *in situ* of low-abundance proteolytic fragments in human brain and animal models.

## 2. Materials and Methods

### 2.1 Antibodies

Primary antibodies used were anti-NeuN, Millipore; anti-GFAP, Abcam; anti-N-terminal C99, 6C3, Abcam, anti-Tau AF3494, R&D, anti-Abeta 6E10, Covance, and RU-369 produced at the Rockefeller University. Secondary antibody was AlexaFluor 488 from Molecular Probes.

### 2.2 Proximity ligation assay (PLA) in cells

N2A cells stably overexpressing APP-695 were cultured in DMEM medium (Gibco, catalog number 11965-092), Opti-MEM (Gibco, catalog number 51985-034), supplemented with 5% FBS (Sigma, catalog number F4135), 100 ug/ml of Penicillin/Streptomycin (Gibco, catalog number 15140-122) and 500 ug/ml Geneticin (Gibco, catalog number 10131-027). Twenty thousand cells were seeded in 12 mm glass coverslips (Corning, catalog number 354085) in a12-well plate and allowed to attach for 24 hrs. Next, cells were incubated overnight with 5 uM BACE inhibitor IV (Millipore Sigma, catalog number 565788) or 1 uM gamma-secretase inhibitor L-685,458 (Sigma, catalog number L1790) or DMSO (Sigma, catalog number 276855) 0.1% as a control.

Next day, the cells were washed with PBS, fixed in 4% paraformaldehyde in PBS for 10 min at room temperature, permeabilized in PBST (PBS with 0.2 % Triton X-100) for 10 min, blocked in 10% goat serum in PBST and incubated in primary antibody overnight in blocking solution. Primary antibodies used to recognize C99 were: RU-369 (Immune serum raised in rabbit, 1:50 dilution) and 6C3 (Abcam, catalog number 126649 raised in mouse, 1:50 dilution). Next, the Duolink kit was used to develop the PLA signal (Sigma, catalog number DUO92008). Briefly, cells were then washed twice for 10 min each time in wash buffer A, incubated with PLA probe solution for 1 h at 37 °C, washed again in wash buffer A, and incubated in ligation-ligase solution for 30 min at 37 °C. Coverslips were washed in wash buffer A and incubated in amplification-polymerase solution for 100 min at 37 °C, washed twice for 10 min each time in wash buffer B. Next, the coverslips were mounted in Duolink mounting media with DAPI. The specificity of the signal was controlled by omitting one of the antibodies. Cells were imaged using a Zeiss LSM710 laser confocal microscope. Quantification of PLA signal was performed using Image J software (NIH).

N2A cells stably overexpressing APP-695 were seeded on sterile, Zeiss high-performance cover glass at a density of 175,000 cells/well and grown overnight. Cells were treated with DMSO or a gamma secretase inhibitor (1 µM) in media for 3 hours. Cells were fixed with 4% PFA in PBS for 10 min at room temperature, permeabilized with PBST (PBS with 0.2 % Triton X-100) for 10 min at room temperature, and blocked in Duolink® PLA blocking solution for 1 hour at 37 °C. The cells were then incubated with primary antibodies RU-369 and 6C3, both at a 1:50 dilution in the Duolink® antibody diluent, overnight at 4°C. PLA was conducted the next day using the Duolink® PLA kit according to manufacturer’s instructions until the mounting step in which the cells were separately stained for DAPI and then mounted onto a glass slide using VECTASHIELD® Antifade Mounting Medium without DAPI (Vector, catalog number H-1000). Coverslips were sealed with metallic nail polish and stored in the dark at 4°C until imaging. Super resolution three-dimensional images were taken using a DeltaVision OMX™ microscope in Structured Illumination mode. 568 nm and 405 nm lasers were used to image DAPI and PLA. Camera temperatures were kept at −80 °C, chamber temperature was kept at 23 °C, and an immersion oil with a refractive index of 1.515 was used. Z-stacks were taken to include all visible PLA dots and nuclei on the coverslip. The image data acquired by DeltaVision OMX was then reconstructed using softWoRx Imaging Workstation. Linespacing, best fit for K0, and amplitude values were evaluated to ensure the quality of the reconstruction. The reconstruction was then subjected to further processing by the “OMX Align Image” function. The final image was then compared to a widefield version of the same image using the “Generate Widefield” and “Deconvolve” functions on the raw non-reconstructed data. Individual sections from the total z-stack were used for comparison.

### 2.3 Proximity ligation assay in human brain sections

Forty nine human brain slides from hippocampus, frontal cortex and occipital cortex were obtained from the brain bank at Mount Sinai/JJ Peters VA Medical Center NIH Brain and Tissue Repository. Donors were selected to evidence no neuropathology, or only the neuropathology of AD excluding any other discernable neuropathology, such a cerebrovascular disease or Lewy bodies. Before proceeding with the staining protocol, the slides were deparaffinized and rehydrated. Slides were placed in a rack, and the following washes were performed: 1. Xylene: 2 × 3 minutes. 2. Xylene 1:1 with 100% ethanol: 3 minutes 100% ethanol: 2 × 3 minutes 4. 95% ethanol: 3 minutes 5. 70 % ethanol: 3 minutes 6. 50 % ethanol: 3 minutes. At the end of the washes, the brain slides were rinsed with water. Antigen retrieval was performed by microwaving the slides for 20 min in citric buffer, pH 6.0. Next, the slides were permeabilized in PBST (PBS with 0.2 % Triton X-100) for 10 min, blocked in 10% goat serum in PBST and incubated in primary antibody overnight in blocking solution. Primary antibodies used to recognize C99 were: RU-369 (Immune serum raised in rabbit, 1:50 dilution) and 6C3 (Abcam, catalog number 126649 raised in mouse, 1:50 dilution). Next, the Duolink kit was used to develop the PLA signal (Sigma, catalog number DUO92008). Cells were then washed twice for 10 min each time in wash buffer A, incubated with PLA probe solution for 1 h at 37 °C, washed again in wash buffer A, and incubated in ligation-ligase solution for 30 min at 37 °C. Coverslips were washed in wash buffer A and incubated in amplification-polymerase solution for 100 min at 37 °C, washed twice for 10 min each time in wash buffer B. Next, the slides were incubated with 0.001% DAPI in 0.01% wash buffer B for 5 min. Then, to quench autofluorescence, the slides were treated with “Trueblack” (Biotum, catalog number 23007) in 70% ethanol for 1 min. and subsequently mounted using Fluoromount-G (Southern Biotech). Slides were imaged using a Zeiss LSM710 laser confocal microscope. For coimmunofluorescence of C99-PLA with neuron and astrocyte markers, as well as with Tau protein, respective antibodies were used together with C99 primary antibodies and secondary antibodies together with PLA probes.

### 2.4 PLA quantification

C99-PLA produces bright red dots that were quantified using Image J software (NIH). The tiff images were thresholded, and the function “Analyze particles” was applied in the red channel. In the blue channel, corresponding to DAPI staining, the function “Cell counter” was applied after threshold.

### 2.5 OMX 3D-SIM using DeltaVision OMX

N2A695 cells were seeded on sterile, Zeiss high-performance cover glass at a density of 175,000 cells/well and grown overnight. Cells were treated with DMSO or a gamma secretase inhibitor (1 µM) in media for 3 hours. Cells were fixed with 4% PFA in PBS and then permeabilized with 0.2% PBS-Triton. Cells were blocked for one hour in Duolink® PLA blocking solution at 37°C. The cells were then incubated with primary antibodies APP369 and C99, both at a 1:50 dilution in the Duolink® antibody diluent, overnight at 4°C. PLA was conducted the next day using the Duolink® PLA kit according to the manufacturer’s instructions until the mounting step in which the cells were separately stained for DAPI and then mounted onto a glass slide using VECTASHIELD® Antifade Mounting Medium without DAPI. Coverslips were sealed with metallic nail polish and stored in the dark at 4°C until imaging. Super resolution three-dimensional images were taken using a DeltaVision OMX™ microscope in Structured Illumination mode. 568 nm and 405 nm lasers were used to image DAPI and PLA. Camera temperatures were kept at −80°, chamber temperature was kept at 23°C, and an immersion oil with a refractive index of 1.515 was used. Z-stacks were taken to include all visible PLA dots and nuclei on the coverslip. The image data acquired by DeltaVision OMX was then reconstructed using softWoRx Imaging Workstation. Linespacing, best fit for K0, and amplitude values were evaluated to ensure the quality of the reconstruction. The reconstruction was then subjected to further processing by the OMX Align Image function. The final image was then compared to a widefield version of the same image using the generate widefield and deconvolve functions on the raw non-reconstructed data. Individual sections from the total z-stack were used for comparisons.

### 2.6 Amyloid plaque staining

Sections from paraffin-embedded blocks were stained using hematoxylin-eosin and anti–β amyloid 4G8 and anti-tau AD2. All neuropathological data regarding the extent and distribution of neuropathological lesions were collected by neuropathologists unaware of any of the clinical and psychometric data.

For quantitative measures of plaque density, 5 representative high-power fields (0.5 mm^2^) were examined and an average density score was calculated for each region and expressed as mean plaque density per square millimeter. When plaques were unevenly distributed in each slide, plaques in the region with the highest density were counted. These sections were adjacent to the sections used to quantify C99 levels.

### 2.7 Statistical analysis

The data are presented as means ± SD. The significance level was determined using a two-way ANOVA with Bonferroni post-test, comparing severe versus non-demented control, moderate versus non-demented control and severe versus moderate. Multiple regression analysis in figure 4B was performed using SPSS statistical package from IBM. Other statistical analyses were performed by using GraphPad Prism software.

## 3. Results

First, we screened several antibodies aimed at recognizing the N and C-terminal domains of C99. The combination of 6C3, a mouse monoclonal antibody [12] which recognizes the cleaved N-terminal of C99 and RU-369, a rabbit polyclonal antibody [13] designed to recognize the C-terminal of APP, successfully recognized C99 (Figure 1A). As expected with a C99-specific marker, the signal was greatly reduced by a BACE inhibitor, which blocks C99 synthesis, and dramatically increased by a gamma-secretase inhibitor, which blocks C99 breakdown (Figure 1B). To further exclude the possibility that the 6C3 antibody recognizes full length APP, we performed double immunofluorescence with 6C3 and RU-369 in N2A-695 cells treated with DMSO, BACE inhibitor or gamma-secretase inhibitor. The signal from 6C3 was reduced upon treatment with a BACE inhibitor and increased upon treatment with gamma-secretase inhibitor, suggesting that 6C3 detects C99 and not APP full length. In addition, Western blot analysis indicated that 6C3 recognized only C99 and not APP full length. (Supplementary Fig. 1). C99-PLA produces bright red dots that can easily be quantified. Super-resolution microscopy revealed that each dot, measuring about 200-400 nm, consisted of a group of 2-3 smaller dots, consistent with the idea of aggregation of C99 molecules (Figure 1C).

**Figure 1.**
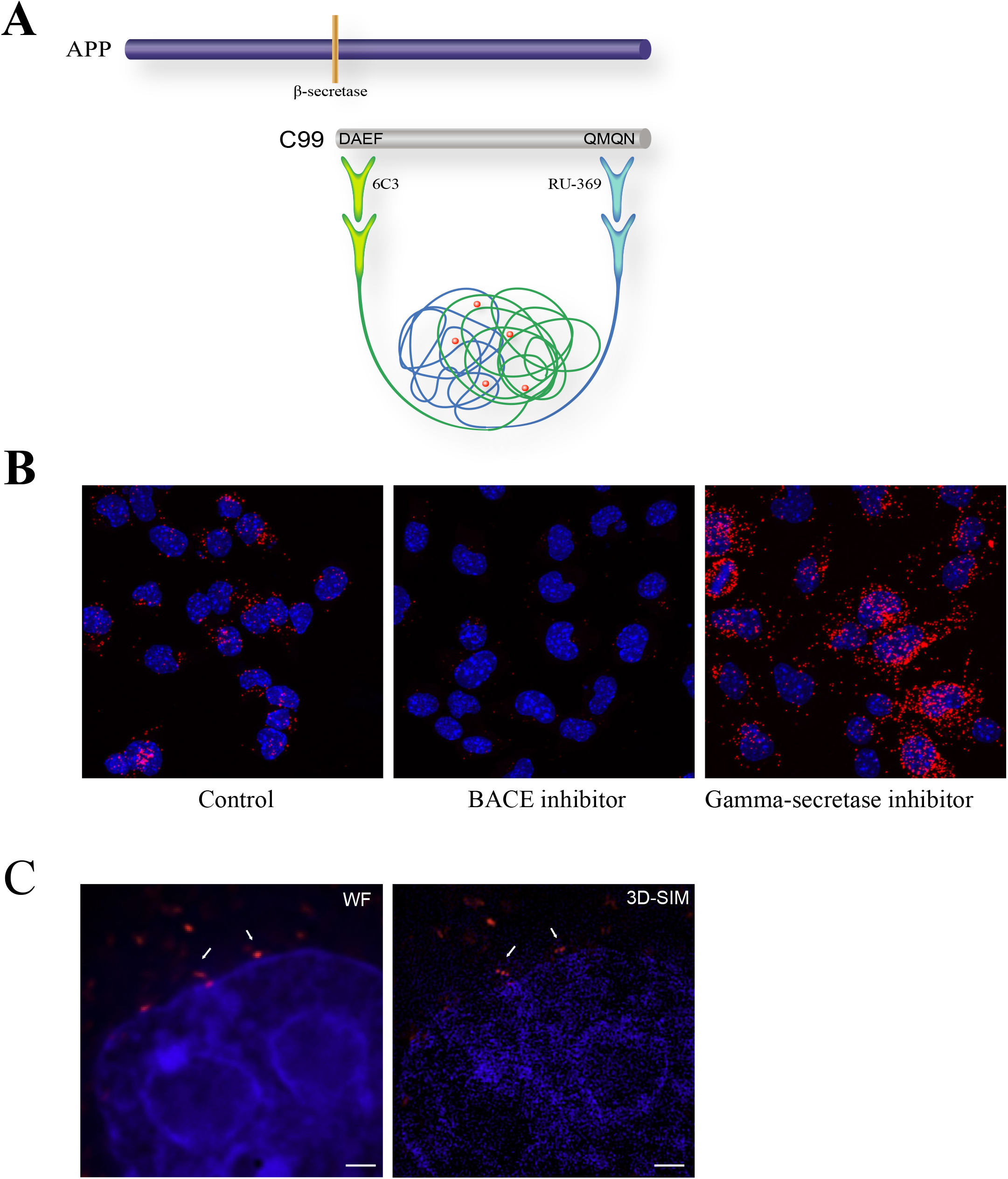
Adaptation of the proximity ligation assay (PLA) for quantitative analysis of C99. A) Diagram showing the location of epitopes for the antibodies (6C3 and RU-369) used in this PLA. B) N2A cells overexpressing APP695 in control conditions (left panel), with BACE inhibitor (middle panel), or with gamma-secretase inhibitor (right panel). Nuclear marker DAPI is in blue and PLA signal is in red. C) Super-resolution microscopy, in wide field (WF, left) and after 3D reconstruction (right), showing that each dot as seen by wide field corresponds to a collection of 2-3 smaller dots (arrows). Scale bar: 1 uM.

Next, we applied this PLA methodology to study the localization of C99 in human brain tissue. Brain tissue sections were obtained from hippocampus, frontal cortex and occipital cortex from 12 controls and 37 patients suffering from sporadic Alzheimer’s disease. Age at the time of death, sex, Braak stage, Thai and CERAD score of these subjects is listed in Supplementary Table 1. PLA specificity was verified by omitting one of the antibodies (Supplementary figure 2). As seen in Figure 2A, C99 is readily detected in the soma and processes of neurons of cognitively normal patients.

**Figure 2.**
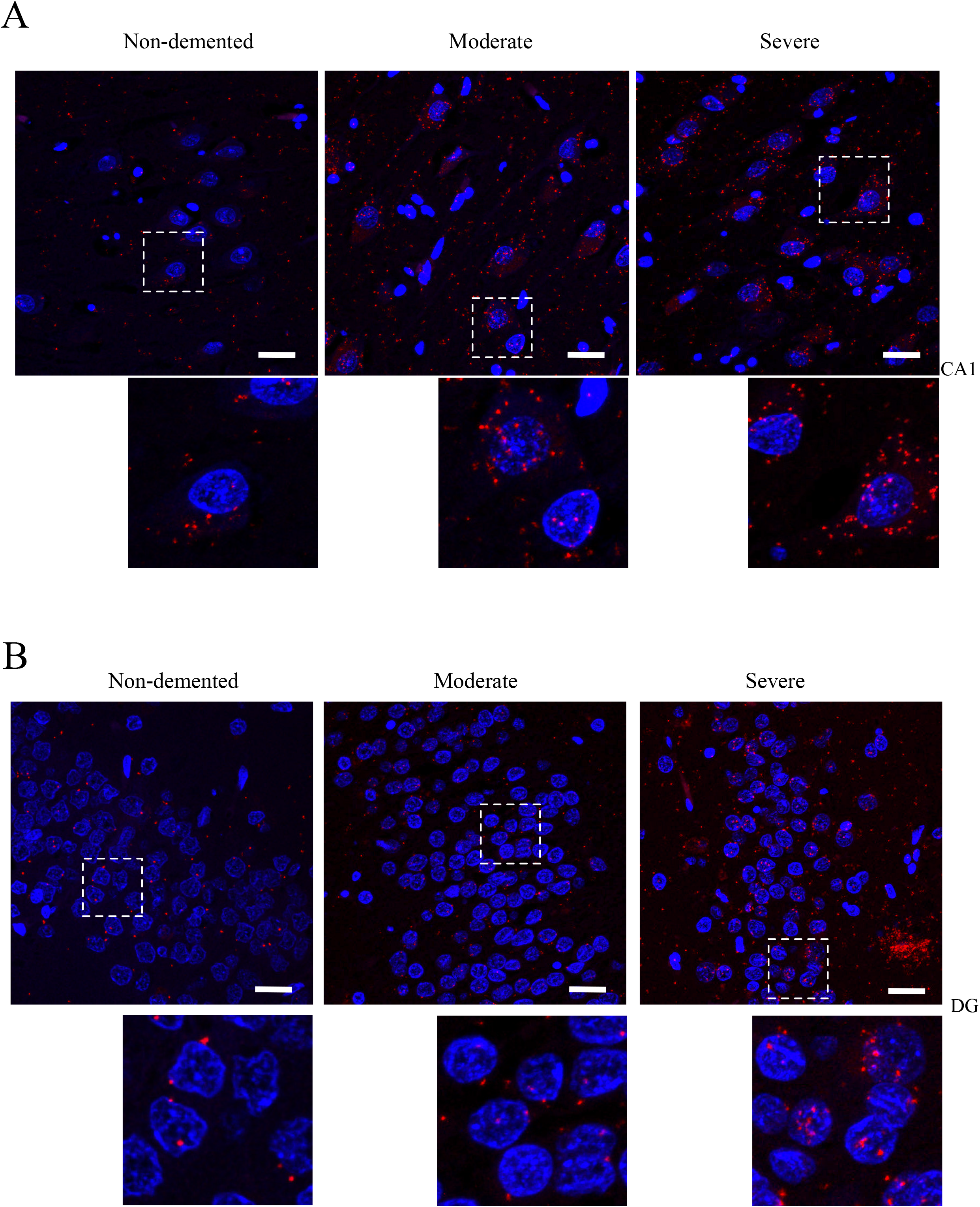
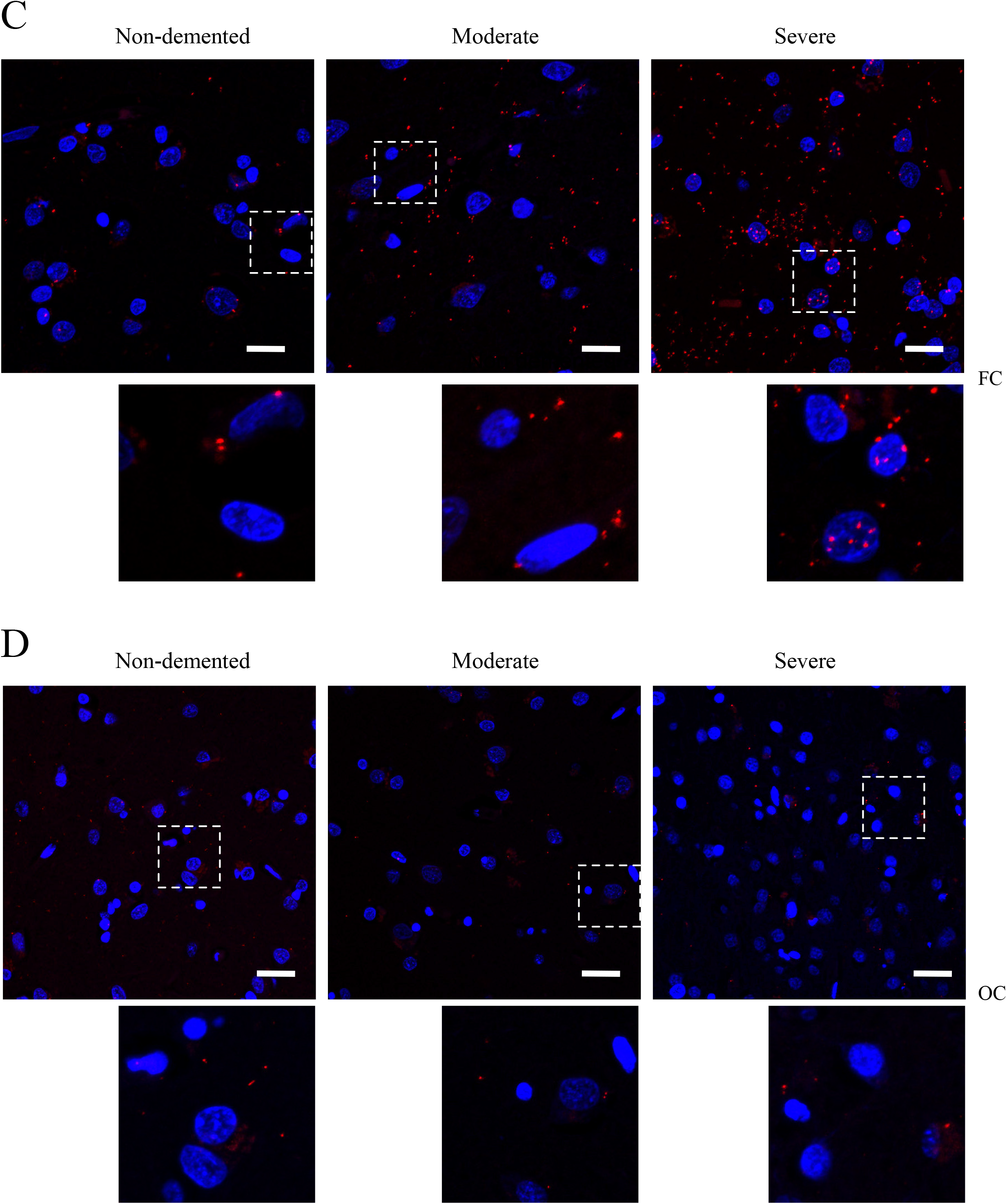
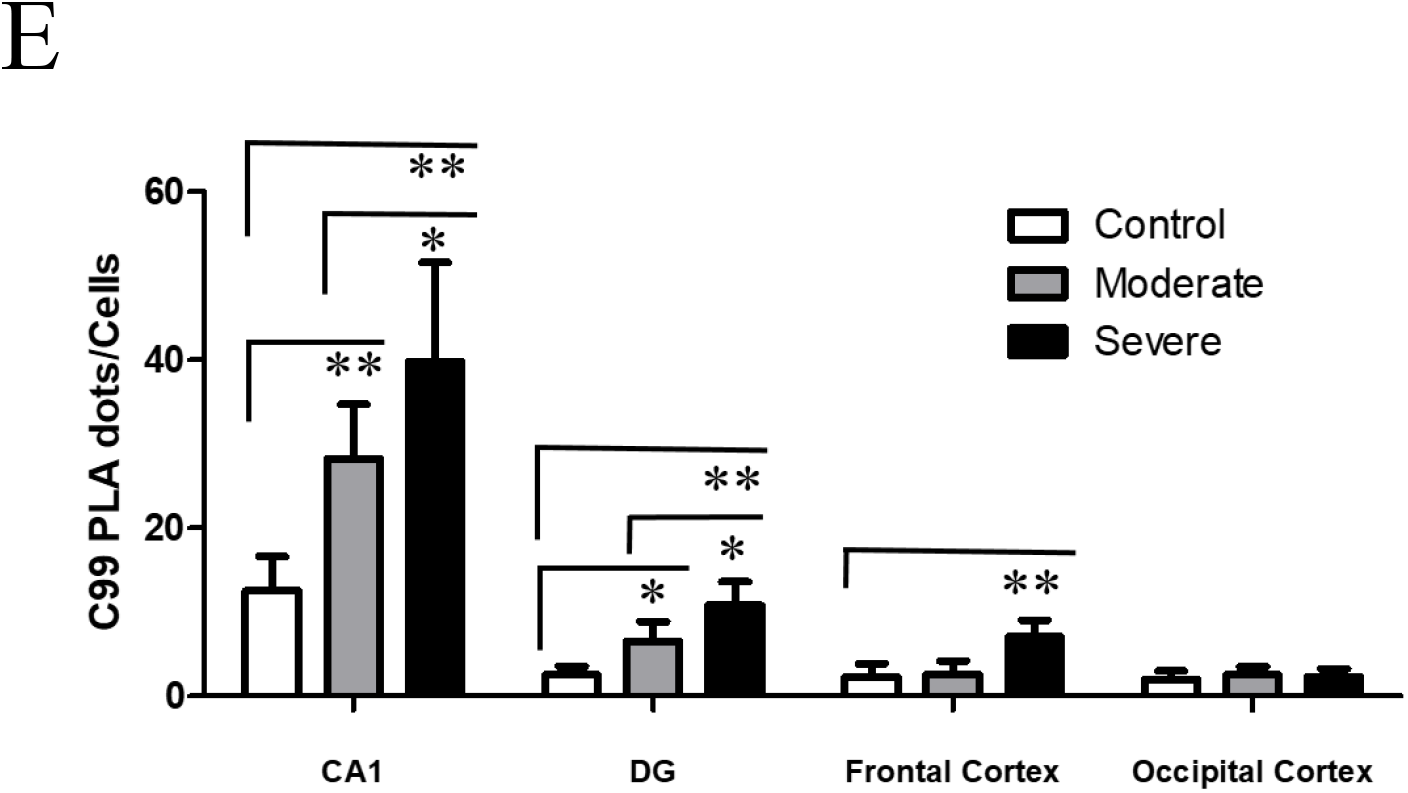
Elevated levels of C99 in vulnerable neurons from brains of AD patients as measured by PLA. Representative images of C99 in A) CA1, B) dentate gyrus (DG), C) frontal cortex (FC) and D) occipital cortex (OC) from moderate and severe Alzheimer’s disease patients and non-demented control subjects. PLA is in red and DAPI in blue. Scale bar: 20 um. E) Quantification of C99 levels in CA1, DG, FC and OC of control, moderate and severe Alzheimer’s disease brains. Data represent means ± SD.* P<0.05, * P<0.01, ANOVA two way with Bonferroni post-test.

To investigate whether C99 levels correlate with cognitive decline, three postmortem groups were studied: subjects with normal and stable cognition, as measured by the clinical dementia rating (CDR = 0), subjects with moderate AD (CDR = 0.5-2), and subjects with severe AD (CDR = 3-5). Average C99 levels were increased in the CA1 area of the hippocampus when comparing severe AD to non-demented controls (39.8 +/− 11.8 vs 12.5 +/− 4.1 (PLA dots/cell), p< 0.001), moderate AD to controls (28.3 +/− 6.4 vs 12.5 +/− 4.1, p<0.001) or severe AD to moderate AD (39.8 +/− 11.8 vs 28.3 +/− 6.4, p<0.05) (Figure 2A, E). In the dentate gyrus area, C99 levels were increased when comparing severe AD to non-demented controls (10.8 +/− 2.8 vs 2.5 +/− 1.1, p<0.001), moderate AD to non-demented controls (6.5 +/− 2.4 vs 2.5 +/− 1.1, p<0.05) and severe AD to moderate AD (10.8 +/− 2.8 vs 6.5 +/− 2.4) (Figure 2B, E). In the middle frontal gyrus of the frontal cortex, C99 levels were increased when comparing severe AD to controls (7.1 +/− 2.0 vs 2.3 +/− 1.6, p<0.001) and severe AD to moderate AD (7.1 +/− 2.0 vs 2.60 +/− 1.6, p<0.001) (Figure 2C, E). Levels of C99 in occipital cortex, an area considered to be resistant to neurodegeneration, were similar in all three groups (Figure 2D, E). Four additional samples from hippocampus of non-demented patients and with severe AD were subject to Western blot analysis using the antibody 6E10. A 4.5 times increase in C99 levels was observed when comparing severe AD to non-demented controls, comparable to the increase seen by PLA (Supplementary figure 3)

**Fig. 3.**
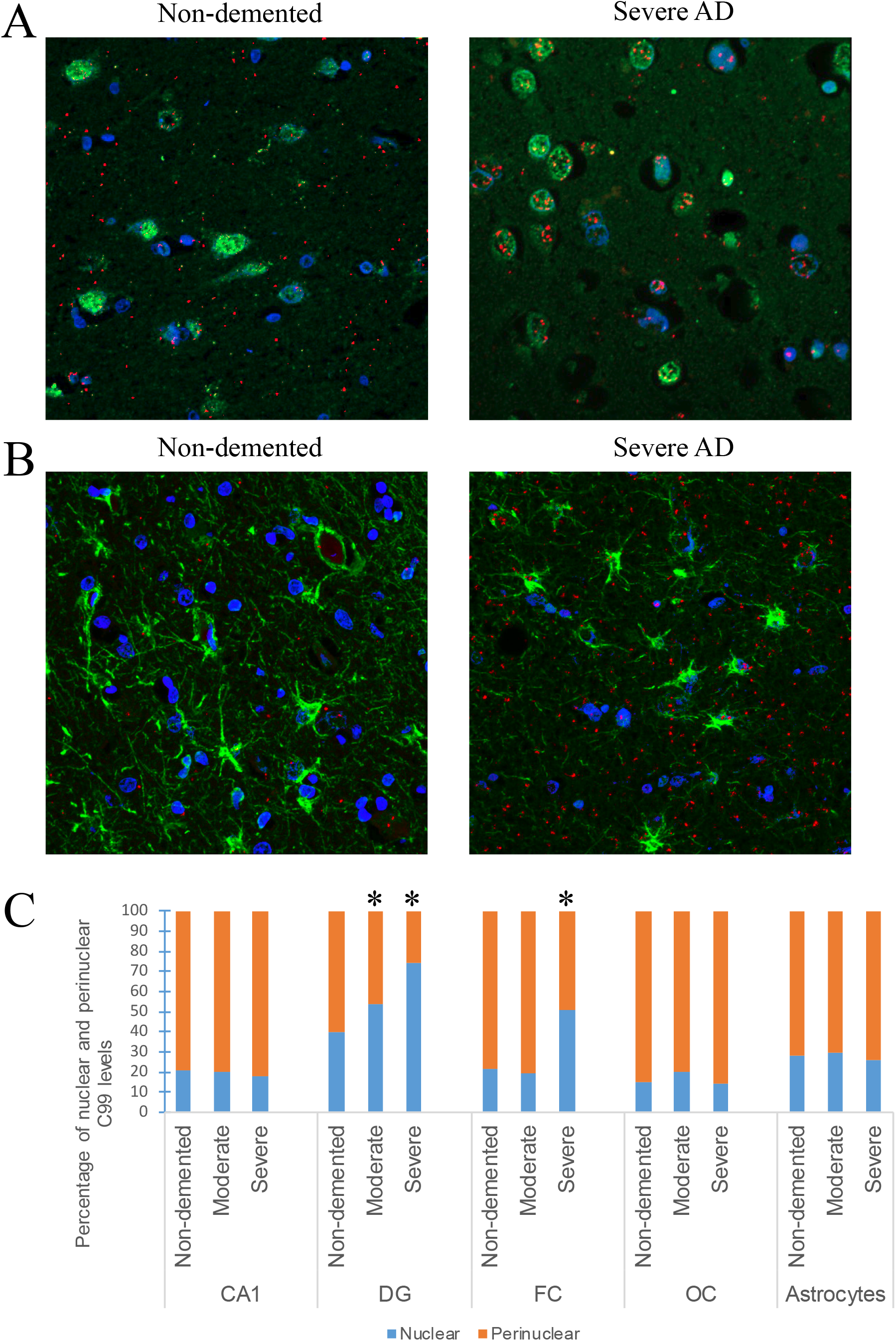
Aberrant localization of C99 in vulnerable neurons in AD. A) C99 distribution in neurons in frontal cortex from a non-demented control subject (left panel) compared to a severe case of AD (right panel). Note the colocalization with NeuN in AD. PLA is in red, NeuN is in green and DAPI in blue. B) C99 distribution in astrocytes from a non-demented control subject (left panel) compared to a severe case of AD (right panel). C) Quantification of the proportion of nuclear and perinuclear C99 label. * P<0.05, ANOVA with Bonferroni post-test comparing against non-demented subjects by brain area or cell type.

**Fig. 4.**
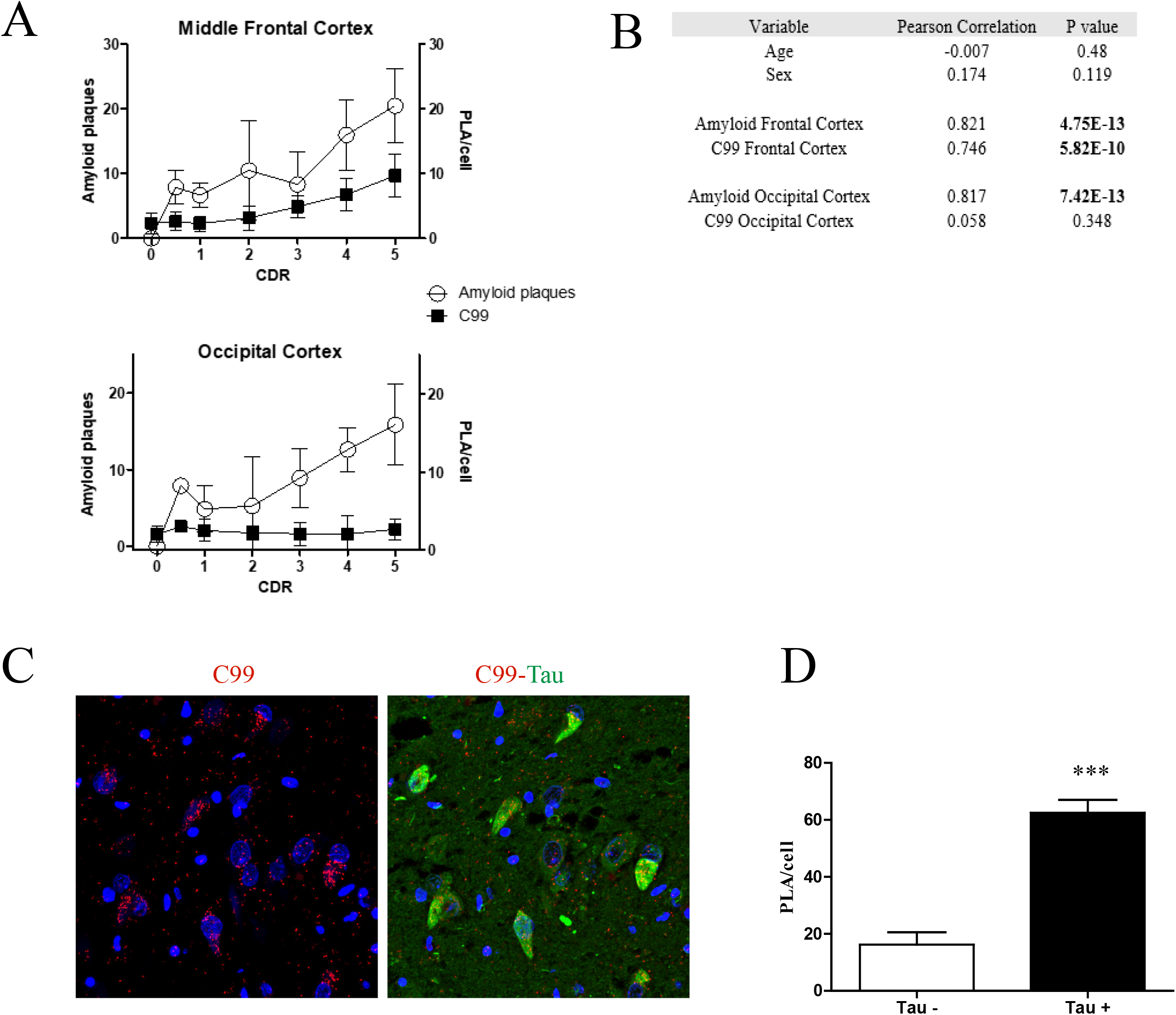
C99 correlates with vulnerability to AD. A) Correlation between cognitive decline and beta-amyloid or C99 levels in the middle frontal cortex (upper graph) or the occipital cortex (lower graph). B) Statistical analysis of the correlation, two-tailed Pearson correlation. C) Left panel: C99-PLA, in red, in CA1 area of moderate AD patient. Right panel: Co-staining with total Tau, in green. Note all the Tau+ neurons contain high levels of C99. D) Quantification of C99 levels in Tau- and Tau+ neurons. Data represents means ± SD.* P<0.001, t-test.

Co-staining with the neuronal nuclear marker NeuN showed that C99 accumulated in the perinuclear region of neurons in non-demented subjects. Its distribution changed to an intranuclear pattern in Alzheimer’s disease subjects in dentate gyrus and frontal cortex (Figure 3A **and** 3C). Interestingly, C99 also accumulated in astrocytes in AD subjects, where it was localized in both the soma and processes, as revealed by GFAP staining. However, in contrast to neurons, the distribution of C99 in astrocytes did not change between non-demented and demented subjects (Figure 3B **and** 3C). Next, we compared the levels of beta-amyloid plaques and C99 in middle frontal cortex and occipital cortex. As seen in Figure 4A **and** 4B, beta-amyloid plaques were present in both vulnerable and non-vulnerable areas, while C99 accumulation correlated with disease progression only in the middle frontal cortex, a vulnerable area, but not in the occipital cortex, a resistant area. To study the association of C99 with selective neuronal vulnerability, costaining with Tau was performed in CA1 area of 7 non-demented controls and 15 AD patients. As seen in Figure 4C **and** 4D, C99 levels were greatly increased in neurons with high levels of Tau, which is a widely accepted marker of neuronal vulnerability. These results show that C99 accumulation, as opposed to beta-amyloid plaques, is specific to neurons most vulnerable to neurodegeneration in Alzheimer’s disease.

## 4. Discussion

Alzheimer’s disease is characterized by accumulation of amyloid plaques and paired helical filaments. Mutations that cause early onset AD are located in the genes that code for the amyloid precursor protein, APP, and the enzyme responsible for amyloid production, Presenilin 1/Presenilin 2. These observations have led to the widely-held belief that beta-amyloid has a causative role in the development of AD. However, the distribution of amyloid deposits in the brain does not correlate well with disease progression. These results suggest the possibility that APP metabolites other than beta-amyloid might contribute to Alzheimer’s disease. It has been proposed that C99, a specific APP metabolite, may cause neurodegeneration by inducing endosome enlargement [14, 15], leading to endosomal dysfunction and neuronal death [16-19].

In this study, we have developed a method to detect C99 *in situ*, and applied this method to measure the levels of C99 in postmortem human brain samples of individuals at different stages of Alzheimer’s disease dementia. We found a correlation between levels of C99 and levels of cognitive decline in areas vulnerable to neurodegeneration, such as the hippocampus and frontal cortex. In the hippocampus, we found increased levels of C99 in the CA1 area and dentate gyrus in both moderate and severe AD, whereas in frontal cortex we found increased levels of C99 only in severe cases of AD, consistent with the well-established delayed involvement of frontal cortex in AD pathology. This correlation was absent in areas resistant to neurodegeneration, such as occipital cortex. In contrast to the area-specific increase in C99 levels, all areas studied in AD patients presented abundant amyloid plaques. Further work is required to understand the underlying mechanism by which C99 accumulates in vulnerable areas of the brain. C99 may accumulate as a consequence of increased synthesis or decreased degradation. Several studies show elevated BACE activity in sporadic AD [20-22], which could account for elevation of C99 levels due to increased synthesis. On the other hand, others and we have shown that C99 is degraded through autophagy and the endosomal-lysosomal pathway [23-27], failure of which could account for a reduced degradation. Elucidation of the detailed molecular pathway that leads to the synthesis and breakdown of C99 is likely to reveal targets responsible for the regulation of this metabolite. Such information may greatly accelerate the development of therapeutic agents for the treatment of AD.

One limitation of the method described here is that it recognizes only one type of APP b-C-terminal fragment product of BACE cleavage. BACE 1 cleaves APP between Met671 and Asp672 to generate the carboxyl-terminal fragment C99. BACE1 also cleaves APP between Thr681 and Gln682, yielding C89 [28]. Our method is specific for C99, and is not able to detect C89. Regarding beta-amyloid levels, the method used detects beta-amyloid plaques, and it is possible that selected beta-amyloid species may correlate with neuronal vulnerability. Another limitation of the present study is the relatively small sample size, which allowed us to draw correlations between C99 and amyloid levels with cognitive decline, but prevented us from evaluating the correlation of C99 levels with Braak staging.

Using a new method to detect specific APP metabolites, we show that C99 accumulates selectively in vulnerable neurons, but not in resistant neurons, in the brains of Alzheimer’s disease patients. Approaches to reduce amyloid burden, such as gamma-secretase inhibitors or immunotherapy, despite being successful at reducing beta-amyloid levels, have failed to stop or reduce the cognitive impairment that characterizes Alzheimer’s disease. Semagacestat, a gamma-secretase inhibitor, worsened the symptoms of dementia in a large phase III clinical trial [29]. It was speculated that this action might have been due to inhibiting the cleavage by gamma-secretase of other substrates such as Notch. We suggest here that the clinical failure may have been due, at least in part, to an accumulation of C99.

The results shown here add to growing evidence that C99 accumulation in vulnerable neurons contributes to neuronal death in AD [14, 15, 17, 18, 30-33]. The question arises as to how C99 induces neuronal death. Dysfunction of the endo-lysosomal and autophagic pathways are one of the earliest key events in AD, progressing to widespread failure of intraneuronal waste clearance, leading to neuritic dystrophy and finally neuronal cell death. It has been shown that elevated levels of C99 in fibroblasts from Down syndrome and AD induce endosomal dysfunction by increasing the recruitment of APPL1 to rab5 endosomes, where it stabilizes active GTP-rab5, leading to accelerated endocytosis, endosome swelling and impaired axonal transport of rab5 endosomes [14, 15]. It will be interesting to see the specific sites of C99 accumulation in human brain (for instance, using PLA and various vesicle markers) and the effect of such accumulation on the endo-lysosomal and autophagic pathway in human neurons in culture. It has also been shown that C99 accumulation in the mouse brain leads to an increase in Cathepsin B and Lamp1 positive vesicles, as well as increased levels of LC3II and autophagic dysfunction [17]. Although the exact molecular mechanism by which C99 exert this effects is not yet known, it is becoming clear that C99 accumulation contributes to neuronal dysfunction and ultimately neuronal death.

Our results show a correlation between C99 levels and cognitive impairment. It has been previously found that synapsis loss is the major correlate of cognitive impairment [4]. To analyze the relationship of C99 with synaptic loss, it could be possible to measure synaptic markers like SNAP25 and synaptophysin [34], in both vulnerable and resistant brain areas and correlate them with C99 levels.

The method that we developed to detect specific APP fragments in fixed brain slides can be modified to quantify APP metabolites in human fluids. To that end, it could be possible to quantify C99 levels in plasma and CSF and correlate them with cognitive deterioration and vulnerability to neurodegeneration, with the goal of developing C99 as an AD biomarker.

In conclusion, our results raise the possibility that C99, rather than beta-amyloid, is responsible for the death of nerve cells in Alzheimer’s disease. In that case, beta-secretase inhibitors, which lower both C99 and beta-amyloid, are preferable to gamma-secretase inhibitors, which lower beta-amyloid but increase C99 levels.

## Acknowledgments

We thank Dr. Fred Gorelick and Dr. Yotam Sagi for discussion and Dr. Jean-Pierre Roussarie for providing reagents and discussion. We thank Alison North and the Bio-imaging facility at The Rockefeller University for training and advice in super-resolution microscopy. We thank Elisabeth Griggs for figure composition. This work was supported by the Fisher Center for Alzheimer’s Research Foundation, The Cure Alzheimer’s fund and JPB Grant 794 (to P.G.). V.B. was supported by DOD/USAMRAA Grant W81XWH-14-1-0045.

## Author contributions

M.P. performed the PLA experiments and confocal microscopy imaging. M.H. performed the super-resolution microscopy. V.H. contributed reagents and measured beta-amyloid plaques. M.P., P.G. and V.B. analyzed data. M.P., P.G. and V.B. designed the project and wrote the manuscript.

## Data and materials availability

All data are available in the main text or the supplementary materials.

## Supplementary Materials

**Supplementary Table 1.** Characteristics of the postmortem samples. Severity of disease was characterized by the clinical dementia rating (CDR). Non-demented subjects had a CDR=0. Moderate subject had a CDR=0.5-2. Severe subjects had a CDR=3-5. CERAD score is as follows: Non pathology: CERAD1; Definite AD=CERAD 2; Probable AD= CERAD 3; Possible AD = CERAD 4

| Patient | Diagnose | CDRScore | Age | Sex | Braak and Braak | Thal | CERAD |
| --- | --- | --- | --- | --- | --- | --- | --- |
| 1 | No pathology | 0 | 85 | M | 1 | A1 | 1 |
| 2 | No pathology | 0 | 74 | F | 1 | A1 | 1 |
| 3 | No pathology | 0 | 73 | F | 1 | A1 | 1 |
| 4 | No pathology | 0 | 95 | M | 1 | A1 | 1 |
| 5 | No pathology | 0 | 70 | M | 1 | A1 | 1 |
| 6 | No pathology | 0 | 94 | F | 1 | A0 | 1 |
| 7 | No pathology | 0 | 75 | F | 1 | A0 | 1 |
| 8 | No pathology | 0 | 70 | M | 1 | A0 | 1 |
| 9 | No pathology | 0 | 91 | F | 1 | A0 | 1 |
| 10 | No pathology | 0 | 72 | M | 1 | A0 | 1 |
| 11 | No pathology | 0 | 66 | M | 1 | A0 | 1 |
| 12 | No pathology | 0 | 101 | F | 1 | A0 | 1 |
| 13 | Moderate | 0.5 | 86 | F | 1 | A1 | 4 |
| 14 | Moderate | 0.5 | 97 | F | 5 | A1 | 2 |
| 15 | Moderate | 1 | 96 | F | 1 | A1 | 4 |
| 16 | Moderate | 1 | 87 | M | 4 | A1 | 3 |
| 17 | Moderate | 1 | 93 | M | 1 | A1 | 3 |
| 18 | Moderate | 1 | 95 | F | 4 | A1 | 2 |
| 19 | Moderate | 1 | 84 | M | 6 | A1 | 2 |
| 20 | Moderate | 1 | 92 | F | 5 | A1 | 2 |
| 21 | Moderate | 1 | 81 | M | 3 | A1 | 2 |
| 22 | Moderate | 1 | 86 | M | 4 | A1 | 3 |
| 23 | Moderate | 1 | 68 | M | 6 | A1 | 2 |
| 24 | Moderate | 1 | 84 | M | 3 | A1 | 3 |
| 25 | Moderate | 2 | 86 | M | 4 | A3 | 3 |
| 26 | Moderate | 2 | 83 | M | 3 | A1 | 2 |
| 27 | Moderate | 2 | 73 | M | 6 | A3 | 2 |
| 28 | Moderate | 2 | 90 | F | 6 | A3 | 3 |
| 29 | Moderate | 2 | 88 | M | 2 | A1 | 3 |
| 30 | Moderate | 2 | 92 | M | 3 | A3 | 2 |
| 31 | Severe | 3 | 94 | F | 6 | A3 | 2 |
| 32 | Severe | 3 | 95 | F | 6 | A3 | 3 |
| 33 | Severe | 3 | 99 | F | 6 | A3 | 3 |
| 34 | Severe | 3 | 73 | M | 6 | A3 | 2 |
| 35 | Severe | 3 | 90 | F | 3 | A1 | 3 |
| 36 | Severe | 4 | 80 | M | 6 | A1 | 2 |
| 37 | Severe | 4 | 97 | F | 6 | A1 | 2 |
| 38 | Severe | 4 | 87 | F | 6 | A1 | 2 |
| 39 | Severe | 4 | 93 | F | 6 | A1 | 2 |
| 40 | Severe | 4 | 81 | F | 6 | A1 | 2 |
| 41 | Severe | 4 | 84 | F | 6 | A3 | 2 |
| 42 | Severe | 5 | 80 | F | 6 | A1 | 2 |
| 43 | Severe | 5 | 80 | M | 5 | A3 | 2 |
| 44 | Severe | 5 | 80 | M | 6 | A3 | 2 |
| 45 | Severe | 5 | 87 | M | 6 | A1 | 2 |
| 46 | Severe | 5 | 65 | F | 6 | A1 | 2 |
| 47 | Severe | 5 | 86 | F | 6 | A1 | 2 |
| 48 | Severe | 5 | 81 | M | 6 | A1 | 2 |
| 49 | Severe | 5 | 73 | F | 6 | A1 | 2 |

**Supplementary figure 1:**
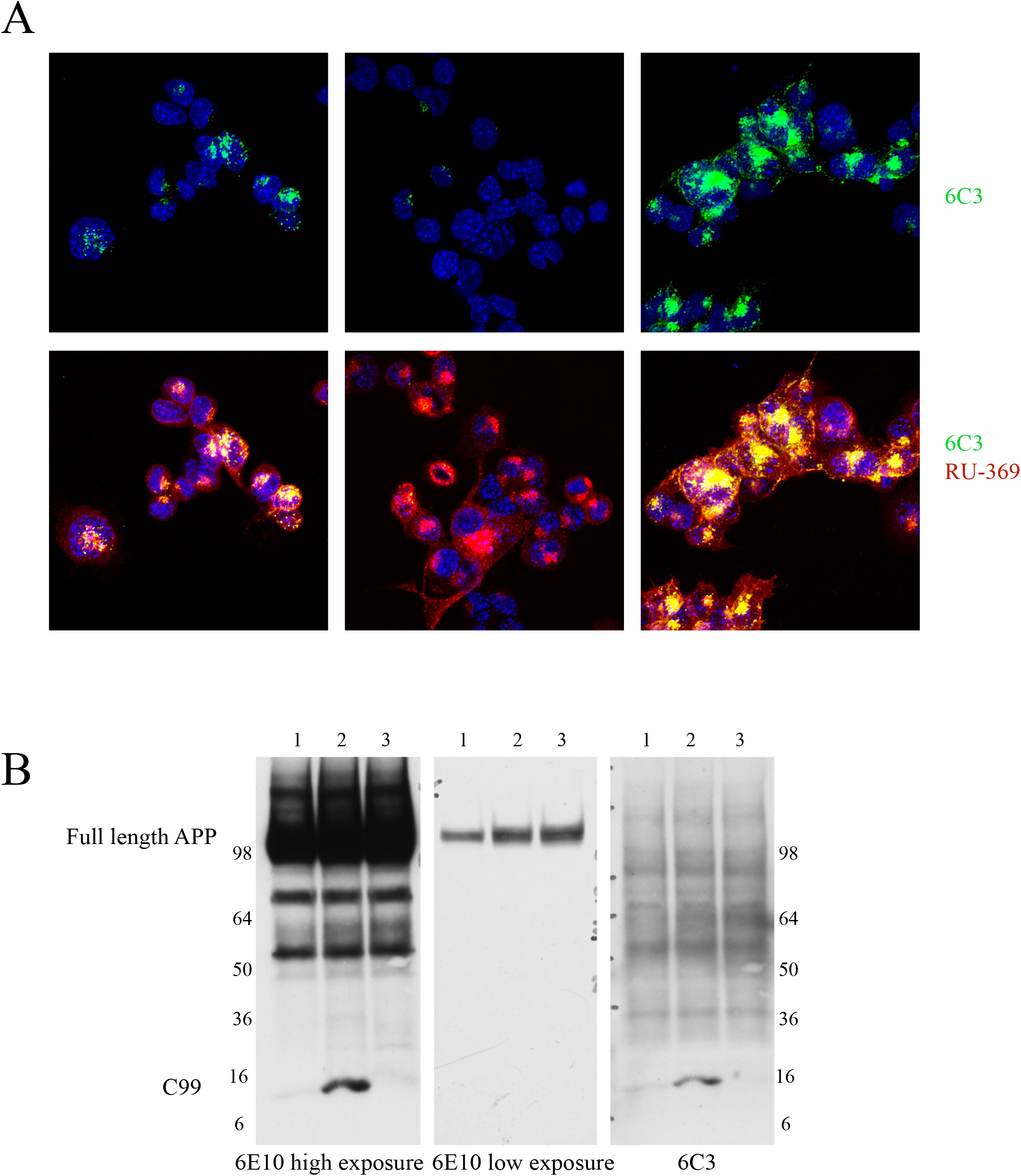
A) Double immunofluorescence with 6C3 (green) and RU-369 (red) of N2A-695 cells treated with DMSO, BACE inhibitor or gamma-secretase inhibitor. Note the signal from 6C3 is greatly reduced upon treatment with a BACE inhibitor and greatly increased upon treatment with gamma-secretase inhibitor, suggesting 6C3 detects C99. B) Western blot with N2A-695 cells treated with DMSO (1), gamma-secretase inhibitor (2) or BACE inhibitor (3) and blotted with 6C3, stripped and reprobed with 6E10 antibody. Note that 6C3, in contrast to 6E10, does not recognizes full length APP.

**Supplementary figure 2.**
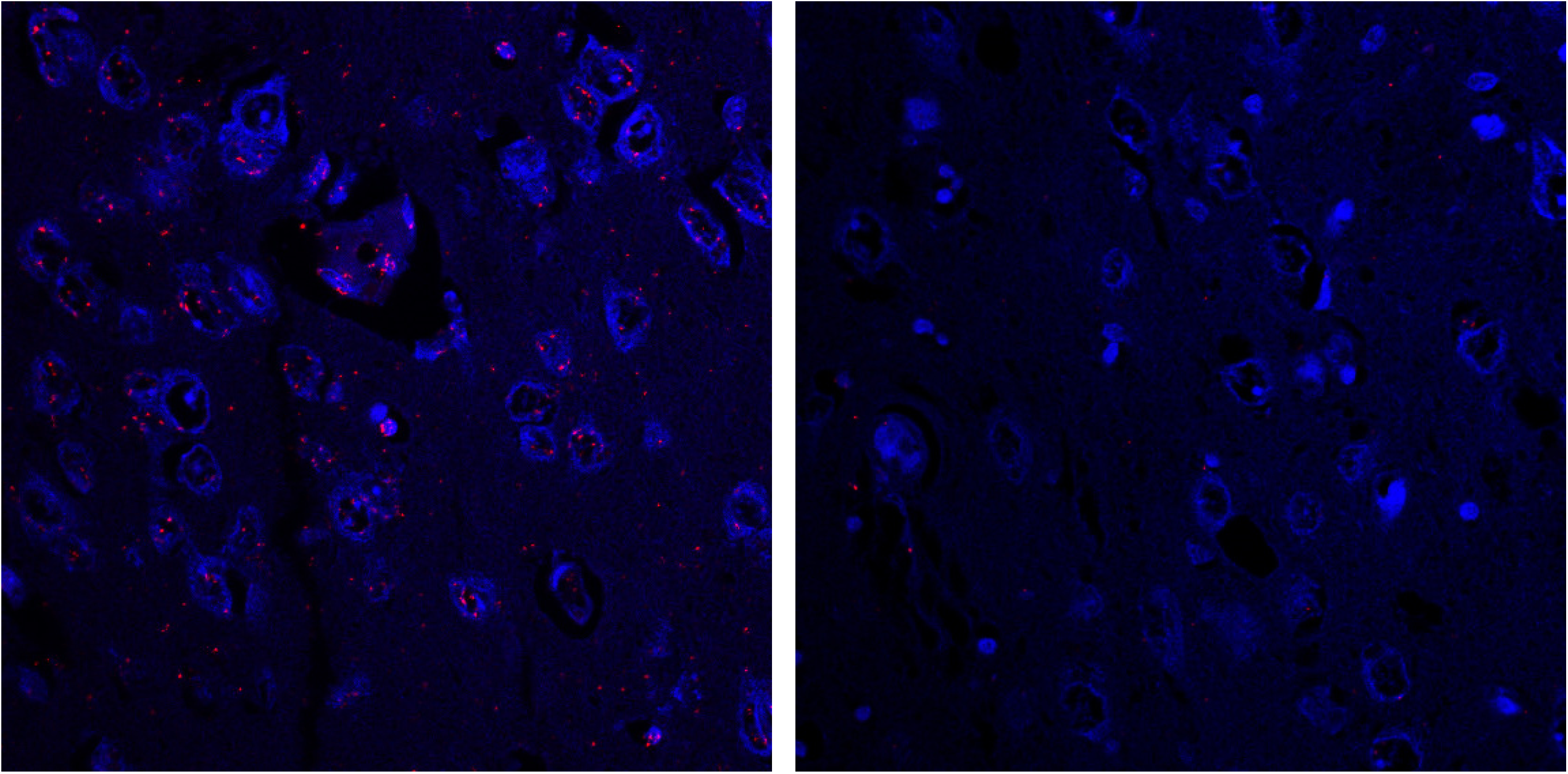
Representative confocal microscopy image of C99-PLA red fluorescent signal (left). Control, where the RU-369 antibody was replaced by rabbit IgG (right), in hippocampus of a patient suffering from Alzheimer’s disease.

**Supplementary figure 3:**
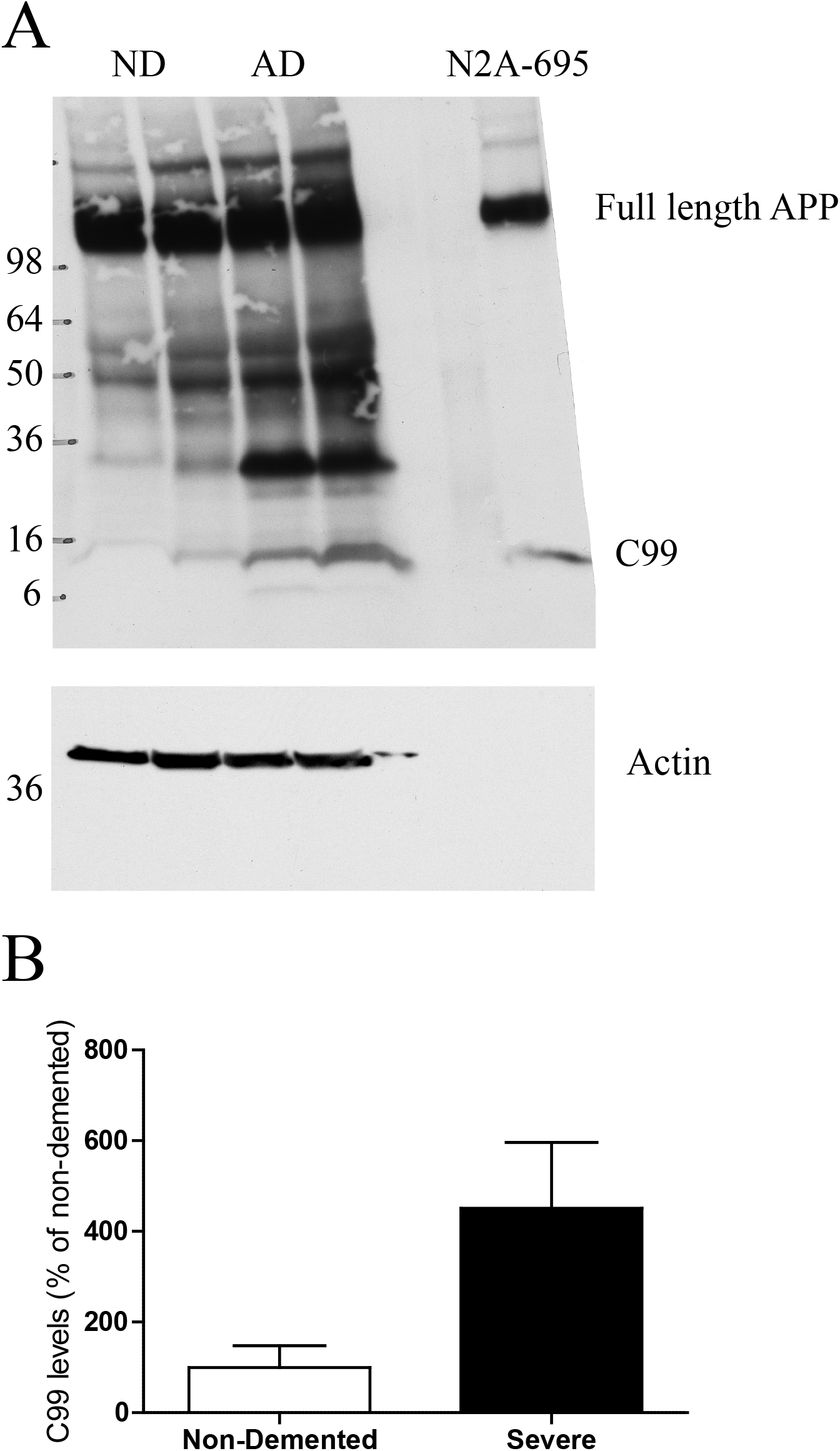
A) Upper blot: Western blot using 6E10 of hippocampal samples from two non-demented (ND) subjects and two severely demented patients (AD). A sample of N2A-695 cells treated with gamma-secretase inhibitor to increase C99 levels was used as a control of size. Lower blot: The blot from the upper panel was stripped and incubated with an antibody against actin, as a loading control. Actin from N2A-695 lysate does not show in this exposure since lower amounts of N2A-695 lysate were loaded to observe the C99 band. B) Quantification of C99 bands from samples on A. A 4.5 times increase in C99 levels was observed when comparing AD versus non-demented samples.

